# The role of 3’UTR-protein complexes in the regulation of protein multifunctionality and subcellular localization

**DOI:** 10.1101/784702

**Authors:** Diogo M. Ribeiro, Alexis Prod’homme, Adrien Teixeira, Andreas Zanzoni, Christine Brun

**Affiliations:** Aix-Marseille University, Inserm, TAGC, UMR_S1090, Marseille, France; CNRS, Marseille, France

## Abstract

Multifunctional proteins often perform their different functions when localized in different subcellular compartments. However, the mechanisms leading to their localization are largely unknown. Recently, 3’UTRs were found to regulate the cellular localization of newly synthesized proteins through the co-translational formation of 3’UTR-protein complexes. Here, we investigate the formation of 3’UTR-protein complexes involving multifunctional proteins by exploiting large-scale protein-protein and protein-RNA interaction networks. Focusing on 238 human ‘extreme multifunctional’ (EMF) proteins, we predicted 1411 3’UTR-protein complexes involving 128 EMF proteins and evaluated their role in regulating protein cellular localization and multifunctionality. Notably, we find that EMF proteins lacking localization addressing signals, yet present at both the nucleus and cell surface, often form 3’UTR-protein complexes. In addition, they provide EMF proteins with the diversity of interaction partners necessary to their multifunctionality. Archetypal moonlighting proteins are also predicted to form 3’UTR-protein complexes thereby reinforcing our findings. Finally, our results indicate that the formation of 3’UTR-protein complex may be a common phenomenon in human cells, affecting up to 20% of the proteins in the human interactome.

## Introduction

Constructing a complex organism does not require a large number of genes. Rather, organism complexity is provided by the ensemble of all the available functions and their timely regulation. Protein multifunctionality, like alternative splicing, allows cells to make more with less. Among multifunctional proteins, “moonlighting” proteins form a particular subset that performs multiple *unrelated* functions^1, 2^ such as the human aconitase, an enzyme of the tricarboxylic acid cycle (TCA cycle) that also functions as a translation regulator, upon an iron-dependent conformational change^3^. However, for most of the moonlighting proteins, the manner with which their distinct functions can be performed, coordinated and regulated is largely unknown. In some cases, the different functions are associated with a change in their *(i)* structural conformation (as for the aconitase) or oligomeric states^4^, *(ii)* interaction partners, *(iii)* location in tissues or cellular compartments. Indeed, in several cases, the presence of a moonlighting protein in different cellular compartments (*e.g.*, nucleus, cytoplasm, plasma membrane) has been found to be responsible for its change in function^5, 6^. For instance, several intracellular chaperones, cytosolic enzymes involved in glycolysis, enzymes of the TCA cycle, as well as other ‘housekeeping’ proteins, also function as cell surface receptors^6, 7^. Among them, RHAMM/HMMR acts intracellularly as a mitotic-spindle or centrosomal protein in the nucleus of normal cells^8, 9^, whereas extracellularly, it is a hyaluronan-binding protein that partners with the CD44 cell-surface protein to signal and promote cell motility and invasion in tumor cells^10^. Interestingly, the RHAMM protein lacks a membrane-spanning domain or other export signals such as an N-terminal signal peptide, in contrast to many other cell-surface receptors^10^. Similarly, out of a compilation of 30 multi-species moonlighting proteins with different functions intracellularly and on the cell surface, none has been found to contain any signal or motif for membrane or cell surface targeting^6, 7^. This suggests that the subcellular localization of multifunctional proteins that participates in their functional diversity may be regulated by a yet unknown mechanism.

In 2015, a breakthrough work by Berkovitz & Mayr^11^ described a novel mechanism directing protein translocation to the plasma membrane. This mechanism involves the interaction between 3’ untranslated mRNA regions (3’UTRs) and RNA-binding proteins (RBPs) during translation, promoting the formation of a protein complex that interacts with the nascent peptide chain^11–13^. In the case of CD47 — a cell-surface protein involved in a range of cellular processes, including apoptosis, adhesion, migration, and phagocytosis^14^— the relationships between alternative 3’UTRs, protein complex formation and subcellular localization have been deciphered in detail^11^. When translated from an mRNA with a short 3’UTR, the CD47 protein is retained in the endoplasmic reticulum, whereas the protein translated from the mRNA with a long 3’UTR localizes to the plasma membrane, thereby affecting its function. Notably, in this manner alternative 3’UTRs can affect the function of their cognate proteins without recurring to amino acid changes. In the CD47 case, this is achieved upon the formation of a 3’UTR-protein complex mediated by the ELAVL1 RBP (also known as HuR), which recognizes a binding site on the long 3’-UTR that is absent from the short one^11^. The complex also contains a specific CD47 protein partner, SET, responsible for addressing the cognate nascent CD47 protein to the plasma membrane. Finally, it has been recently shown that this 3’UTR-protein complex forms within a newly described membraneless subcellular compartment, the TIGER domain, constituted by TIS granules - accumulating another RBP named TIS11B - located at the Endoplasmic Reticulum surface^15^.

The translocation mechanism involving 3’UTR-protein complex formation has the potential to be a widespread trafficking mechanism for proteins located at the membrane^11^. However, its prevalence is not documented. Moreover, alternative 3’UTRs could play a role in mediating the multifunctionality of proteins, as recently shown for BIRC3^16^, by promoting the formation of different complexes containing distinct interaction partners. There is thus a need to determine whether the formation of 3’UTR-protein complexes is a major contributor to the diversification of protein function.

As the moonlighting functions of proteins have usually been discovered by serendipity, we have previously proposed a computational approach combining protein-protein interaction (PPI) network and Gene Ontology annotation analyses to identify at large-scale moonlighting candidates that we termed “extreme multifunctional” (EMF) proteins^17^. We showed that EMF proteins are characterized as a group by particular features, constituting a signature of extreme multifunctionality^17^. Notably, EMF proteins contain more short linear motifs (SLiMs) than other proteins^17^, these short and conserved sequences mostly located in structurally disordered regions, that can mediate transient interactions and be used as molecular switches between functions^18, 19^.

Here, we aim to determine the role of the 3’UTR-protein complexes in regulating protein cellular localization and multifunctionality. We first established EMF proteins as a model to investigate the regulation of multifunctionality mediated by 3’UTRs and then predicted all 3’UTR-protein complexes plausible to be formed with EMF proteins, using large-scale protein-protein and RBP-3’UTR interaction networks. With this approach, we identified more than a thousand possible 3’UTR-protein complexes, comprising 128 out of 238 EMF proteins. Comprehensive computational analysis of the composition of the predicted 3’UTR-complexes led us to propose that the translocation of proteins between subcellular compartments — particularly to the plasma membrane — which is often associated with the functional change of multifunctional proteins could be mediated by their 3’UTRs. Notably, our hypothesis is largely supported by the finding of numerous well known moonlighting proteins among our predicted 3’UTR-protein complexes. In addition, we extend the current knowledge on the few experimentally described 3’UTR-protein complexes^11, 16, 20^ by predicting that as much as 20% of proteins in the human PPI network (i.e., the interactome) are able to form such complexes. The formation of 3’UTR-protein complexes could, therefore, represent a common protein trafficking mechanism that has been so far largely overlooked and underestimated.

## Results

### EMF proteins as a model to study the regulation of protein localization and multifunctionality by alternative 3’UTRs

The usage of alternative 3’UTRs has been found to regulate the subcellular localization of proteins^11, 21^, which in turn could regulate their functions. Therefore, we hypothesize that 3’UTRs may play a role in moonlighting protein function regulation. To investigate this possibility, we explore a comprehensive set of human moonlighting protein candidates, *i.e.* 238 ‘extreme multifunctional’ (EMF) proteins that we identified using our MoonGO approach^17^ and that are available in our recently updated MoonDB 2.0 database^22^ (see Methods).

Numerous moonlighting proteins have been found to perform different functions when localized in different cellular compartments. Coherently, here we found that EMF proteins are annotated with significantly more “Cellular Component” (CC) GO terms than other groups of proteins (see Methods, Fig. 1a). Indeed, on average an EMF protein is associated with 7.8 CC GO terms, whereas the average in the PPI network — from which the EMF proteins are identified — is 4.6 CC GO terms (Mann-Whitney U test P-value = 1.1 x 10^−22^). We also observed that 111 EMF proteins (46.6% of the total) belong to the multilocalizing proteome, defined by immunofluorescence-based approaches in the Human Protein Atlas (HPA)^23^, a significantly higher fraction than in the PPI network (36.7%, two-sided Fisher’s Exact Test, odds ratio (OR) = 1.52, P-value=1.7 x 10^−3^) (Fig. 1b). Moreover, using a dataset of 1233 known and machine learning-predicted translocating proteins from the Translocatome database^24^, we found that 41.2% EMF proteins (98 out of 238) could change their subcellular localization upon a regulatory event, representing, again, a higher fraction than in the PPI network (9%, two-sided Fisher’s Exact Test, OR = 7.8, P-value < 2.2 x 10^−16^) (Fig. 1b). Overall, this indicates that EMF proteins are more often localized in different cellular districts than expected, a feature that may be regulated by alternative 3’UTRs.

**Figure 1.**
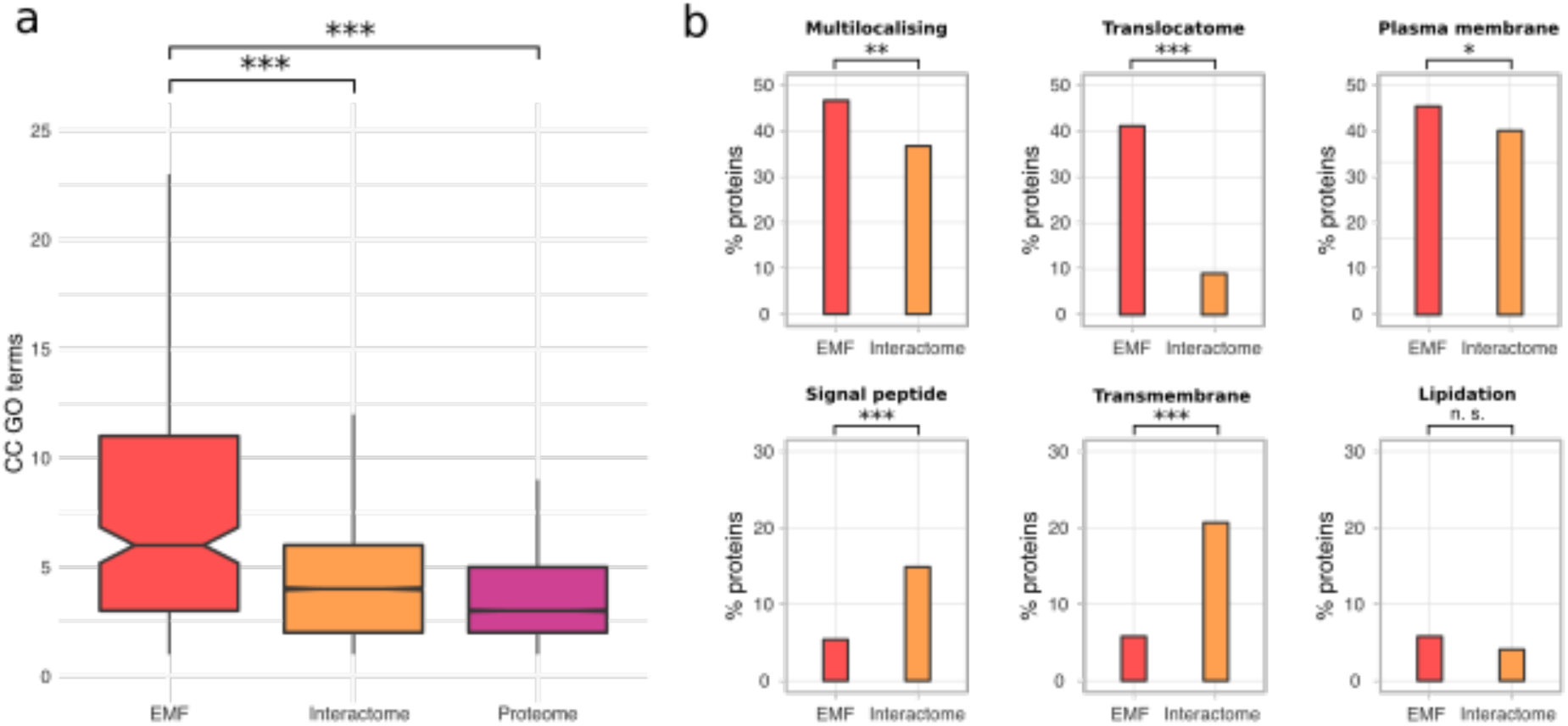
Cellular localization of EMF proteins. **(a)** Comparison of number of Cellular Component (CC) GO term annotations between protein groups (Kruskal-Wallis rank sum test, P-value < 2 x 10^−16^): EMF (n=235), interactome (n=12,794), and proteome (n=18,298). Mann-Whitney U tests were performed to assess pairwise statistical significance. The Benjamini-Hochberg procedure was applied for multiple test corrections. Significance: ‘***’ indicates a FDR < 0.001. **(b)** Enrichment analysis of cellular localization annotations and membrane-targeting signals. Significance: ‘*’ indicates a P-value < 0.05; ‘**’ indicates a P-value < 0.01; ‘***’ indicates a P-value < 0.001.

Next, given the fact that many proteins have been found to perform moonlighting functions when localized at the plasma membrane, as in the case of RHAMM (see Introduction), we sought for EMF proteins associated with the plasma membrane. Indeed, we found that 65 out of 238 (27.3%) EMF proteins have been observed at the plasma membrane according to GO term annotations (“plasma membrane”, GO:0005886; annotations by manual assertion only, see Methods). This is significantly more than expected when considering the interactome as the background (24.1%, two-sided Fisher’s Exact Test, OR = 1.46, P-value = 1.22 × 10^−2^; Fig. 1b). Coherently, the same trend is observed when considering the proteins localized at the “plasma membrane” according to the Human Protein Atlas (see Methods). However, although enriched in proteins localized at the plasma membrane, very few EMF proteins (31, i.e. 13%) contain a signal peptide or a transmembrane domain or a post-translational modification site (*i.e.*, amino acid lipidation site) that could explain their subcellular location. Strikingly, these features are depleted or not significantly enriched in EMF proteins when compared to the PPI network considered as a group (two-sided Fisher’s Exact Test, Signal peptides: 5.4%, OR =0.32, P-value = 4 × 10^−6^; Transmembrane domains: 5.8%, OR= 0.23, P-value = 1 × 10^−10^; Lipidation sites: 5.8%, OR=1.46, P-value = 0.18, Fig 1b). Altogether, this suggests that part of the EMF proteins are not localized at the membrane *via* the classical route.

In order to estimate the potential of EMF proteins to be regulated by the 3’UTR mechanism, we investigated the features of their 3’UTR sequences. Using 3’UTR models from the Ensembl database^25^, we found that mRNAs encoding EMF proteins have significantly longer 3’UTRs than mRNAs encoding all other human proteins as well as those present in the interactome (see Methods, mean ‘EMF’ (1987 nt) versus ‘Proteome’ (1643 nt), Mann-Whitney U test P-value = 1.5 x 10^−3^; mean EMF (1987 nt) versus ‘Interactome’ (1739 nt), Mann-Whitney U test P-value = 3.0 x 10^−2^) (Fig. 2a).

**Figure 2.**
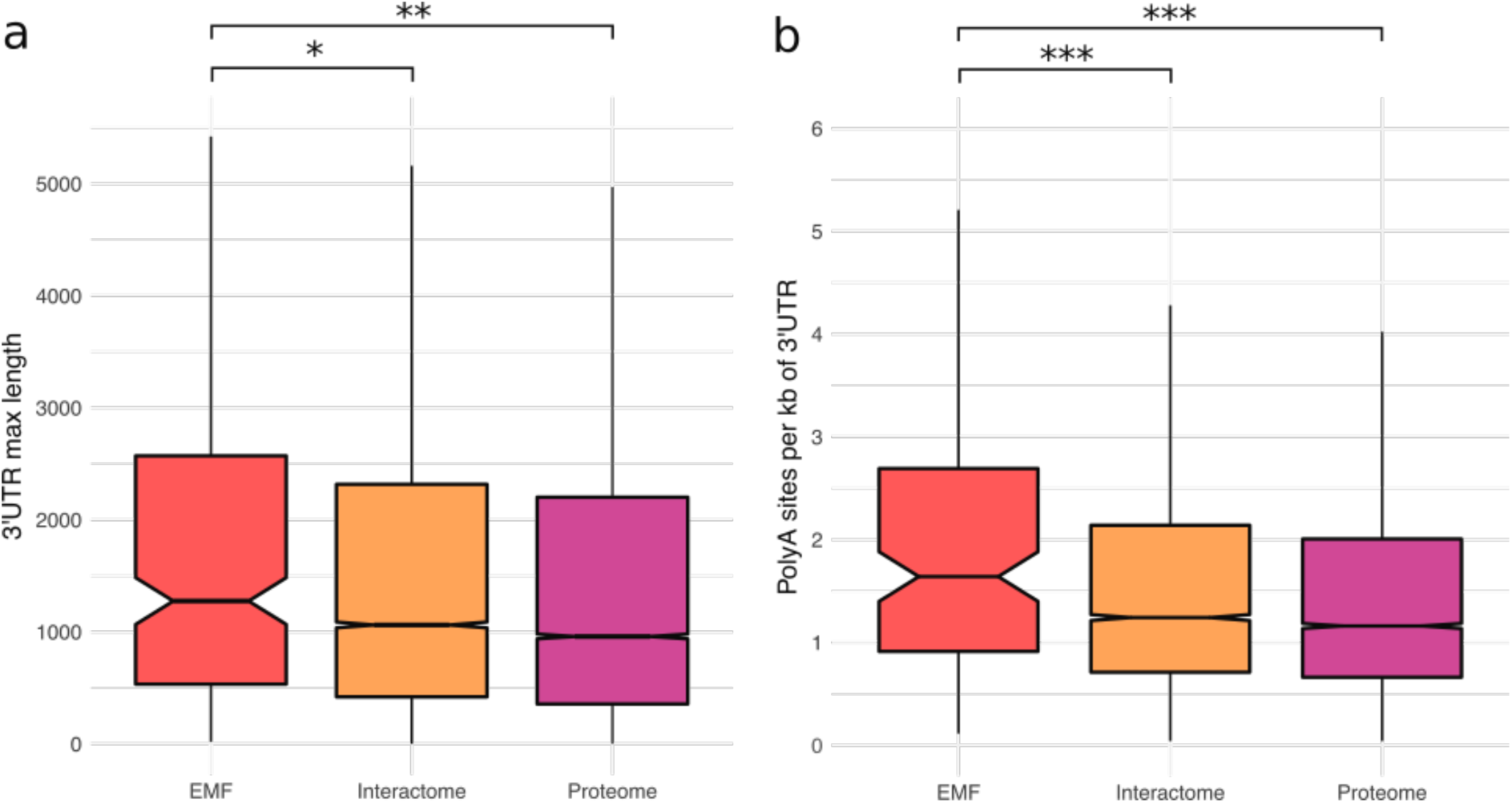
3’UTR-related features of EMF proteins. **(a)** Comparison of maximum 3’UTR lengths. Kruskal-Wallis rank sum test, P-value = 3 x 10^−14^. **(b)** Number of polyadenylation (polyA) sites per kb of 3’UTR, using APADB data^22^. Kruskal-Wallis rank sum test, P-value = 5 x 10^−8^. Only transcripts with 3’UTRs longer than 1000 nucleotides were considered. Mann-Whitney U tests were performed to test for pairwise statistical significance. The Benjamini-Hochberg procedure was applied for multiple test corrections. Significance: ‘*’ indicates a FDR < 0.05; ‘**’ indicates a FDR < 0.01; ‘***’ indicates a FDR < 0.001.

In addition, using polyadenylation sites from APADB^26^ and PolyASite databases^27^, we found that mRNAs encoding EMF proteins bear a higher number of alternative polyadenylation (APA) sites in 3’UTRs than mRNAs encoding the other groups of proteins (Supplementary Fig. 1), even when accounting for the observed differences in 3’UTR length between the protein groups, by calculating the number of APA sites per kb (Fig. 2b; mean EMF (2) versus Interactome (1.6), Mann-Whitney U test P-value = 2.8 x 10^−4^). Consistent with these findings, EMF proteins have significantly more 3’UTR isoforms than the other protein groups (Supplementary Fig. 2). Together, these results suggest that mRNAs encoding EMF proteins are more likely to be regulated by their 3’UTRs than those encoding other proteins.

Overall, the functions of EMF proteins could be affected by their cellular localization, particularly in association with the plasma membrane. Moreover, with longer and more variable 3’UTRs, the EMF proteins thus have the potential to be regulated by a mechanism involving their 3’UTR. EMF proteins, therefore, constitute a suitable model to investigate the role of 3’UTR-protein complex formation in the localization and the multifunctionality of proteins.

### Prediction of 3’UTR-protein complexes

In order to assess the possible involvement of the 3’UTR in the regulation of the localization and the function of EMF proteins, we predicted the potential formation of 3’UTR-protein complexes containing human EMF proteins. As the mechanism involves the recruitment of RBPs to the site of translation by 3’UTRs, which in turn may promote the co-translational formation of protein complexes that interact with the nascent peptide chain^11–13^, the 3’UTR-protein complex formation conceptually involves the following components: (*i)* an mRNA with a 3’UTR; (*ii)* the cognate protein being translated (hereby termed ‘nascent’ protein); (*iii)* an RBP able to bind the 3’UTR; (*iv)* one or more other proteins (hereby termed ‘intermediate’ proteins) that interact with the RBP and the nascent protein. By searching for sets of co-interacting 3’UTRs, RBPs, nascent and intermediate proteins (Fig. 3; see Methods) from two large-scale experimental datasets forming a RBP-3’UTR interaction network (from AURA database^28^) — 163,490 interactions between 201 RBPs and the 3’UTR of the mRNAs of 10893 protein-coding genes — and a PPI network (from MoonDB 2.0^22^) — 90,386 interactions between 14,047 proteins —, we identified possible 3’UTR-protein complexes. To simplify our approach, we only considered the presence of one intermediate protein per complex. Moreover, to identify biologically relevant complexes, we only predicted 3’UTR-protein complexes if the RBP, nascent and intermediate proteins are co-present in at least one of the 58 Human Protein Atlas (HPA) normal tissues^29^.

**Figure 3.**
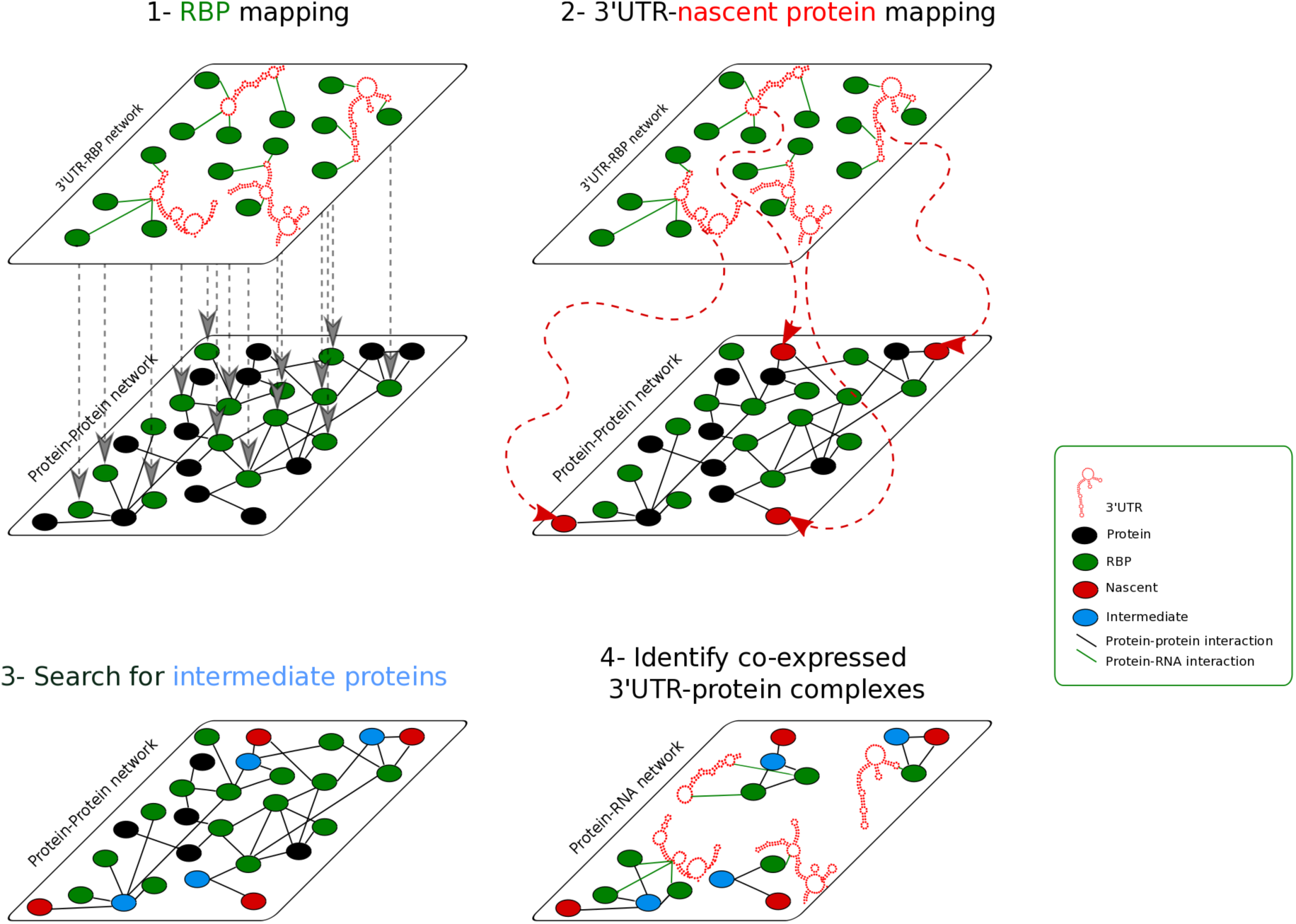
Workflow of the 3’UTR-protein complex prediction. Experimental RBP interactions with 3’UTRs of mRNA are retrieved from AURA v2 database A large-scale PPI network is retrieved from MoonDB 2.0 (see Methods). 3’UTR-protein complexes are predicted by 1-mapping the RBP (green nodes) interacting with 3’UTRs onto the PPI network, 2-finding cases in which the 3’UTR of a ‘nascent’ protein (protein under synthesis, red node) interacts with RBP, which in turn 3-interacts with an interaction partner of the nascent protein (‘intermediate’, blue node). Finally, 4-only 3’UTR complexes where the nascent, RBP and intermediate proteins are present in at least one same tissue are kept (Human Protein Atlas (HPA), 58 normal tissues).

### A set of predicted 3’UTR-complexes containing EMF proteins

Using our novel approach, starting from 238 EMF proteins, we predicted a total of 1411 distinct 3’UTR-protein complexes comprising 128 EMF proteins and a combination of 87 RBPs and 440 interacting intermediate proteins (Table 1, Supplementary Table 1). Notably, 53.8% of the EMF proteins (128 out of 238) may form at least one 3’UTR-protein complex whereas a much lower percentage of the proteins of the interactome do so. Indeed, 16.9% of the proteins of the interactome are found in 9657 3’UTR-complexes (Fisher’s Exact test, two-sided, OR = 6.46, P-value = 2.44 x 10^−43^) (Table 1).

**Table 1.** Summary of the 3’UTR-protein complex prediction. Percentages under the ‘Nascents in complex’ column are relative to the initial number of proteins in the protein group (238 for EMF, 14046 for interactome).

| Protein group | Nascents in complex | RBPs | Intermediates | 3'UTR-protein complexes |
| --- | --- | --- | --- | --- |
| EMF | 128 (53.8%) | 87 | 440 | 1411 |
| Interactome | 2354 (16.8%) | 113 | 789 | 9657 |

To confirm the plausibility of our predictions, we estimate the number of 3’UTR-protein complexes formed by chance while shuffling 10,000 times all proteins in the protein-protein interaction network. We found that the numbers of 3’UTR-protein complexes predicted are 2.8 to 12 times higher than expected by chance for interactome and EMF proteins, respectively (Supplementary Table 2), therefore supporting our predictions.

Only 42% of the RBPs known to interact with a 3’UTR in AURA database, and present in the binary interactome (87 out of 204, Table 1) are found in EMF-containing complexes, despite the general propensity of RBPs to bind a large number of RNAs^30^. This, therefore, increases our confidence in the specificity of the predicted 3’UTR complexes. EMF proteins are often highly connected in the PPI^17^, and thus potentially more likely to form 3’UTR-protein complexes. Indeed, 80% of them are hubs, defined here as nodes whose degree is at least twice the network average (≥32). We, therefore, verified that the number of complexes into which the EMFs participate is poorly correlated with their number of protein partners in the network (Spearman correlation, rho=0.326, P-value=2.66×10^−7^, Supplementary Fig. 3), indicating that the number of interaction partners of EMF proteins do not greatly influence our results.

Alternative 3’UTRs were found to regulate the localization and/or function of CD47^11^ and BIRC3^16^. Notably, we found that the vast majority of the predicted complexes, i.e. 1020 out of 1411 (72,3%),contains EMF proteins with at least two alternative 3’UTR isoforms, (80 EMF proteins out of 128, herein named “mUTR”). Moreover, 60% of these complexes containing mUTR EMF proteins (i.e., 612 out of 1020), are predicted to form exclusively by the binding of RBPs to long isoforms (Fig. 4), as in the known cases. Interestingly, similar results are obtained when we extend the analysis to the 3’UTR-protein complexes predicted on the whole interactome (Supplementary Fig. 4). Altogether, these large-scale results correctly recapitulate the knowledge obtained in the few experimental complexes investigated previously^11, 16^. This, therefore, encouraged us to pursue our analysis and test the possibility that 3’UTR-protein complex formation may contribute to the regulation of EMF – and other protein – functions.

**Figure 4.**
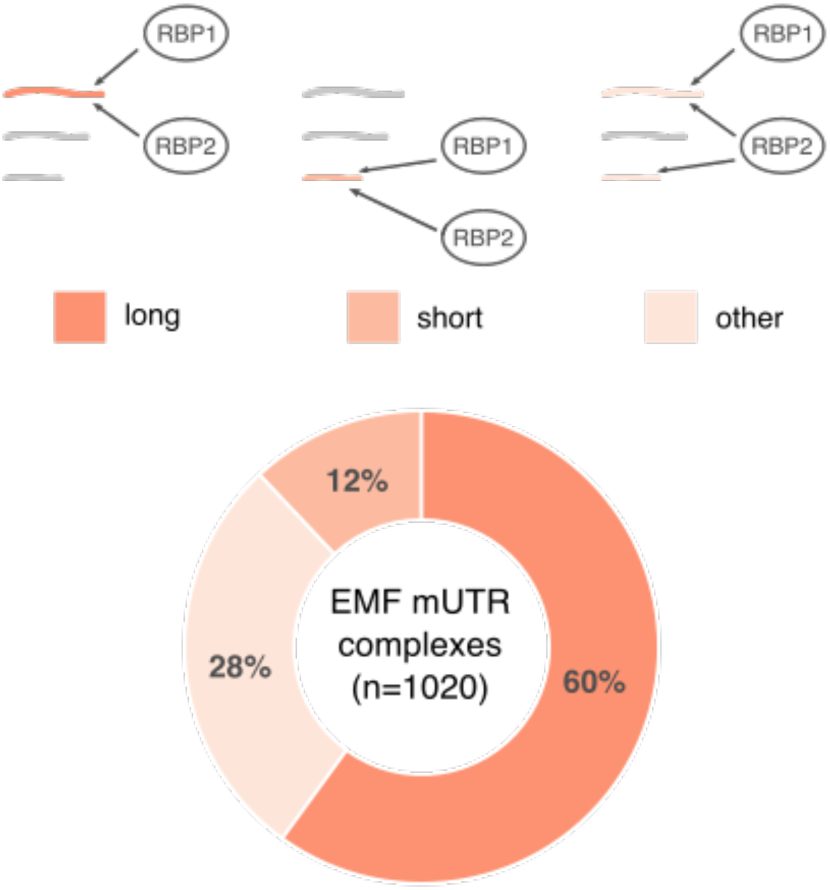
Preferential isoform usage in mUTR EMF-containing predicted complexes. Nascent mUTR EMF proteins were classified according to the length of the 3’UTR(s) bound to RBPs: the longest or the longer isoforms are bound to RBPs (“long”); the shortest or shorter isoforms are bound to RBPs (“short”); RBPs bind all isoforms, or a substantial fraction of them, without a clear preference for long or short isoforms (“other”).

### The role of 3’UTR-protein complexes in the subcellular localization of EMF proteins

The 128 EMF proteins predicted to belong to at least one 3’UTR-protein complex are enriched in plasma membrane annotations. Indeed, according to Gene Ontology annotations, 45 out of the 128 EMF proteins in complexes (i.e. 35%) have been associated to the plasma membrane, a significant enrichment when compared to the human interactome (OR=1.5, two-sided Fisher’s Exact Test, P-value = 3.4 × 10^−2^, Table 2). Remarkably, 36 out of the 45 EMF proteins in complexes and annotated to the plasma membrane (i.e. 80 %) do not display any membrane addressing signal, such as signal peptides, transmembrane or intramembrane domains, or lipid anchors, all features known to be critical for localizing proteins to the membrane^31, 32^. This proportion is remarkably higher than the one in the PPI network, where it drops to 47.5%(OR=4.56, two-sided Fisher’s Exact Test, P-value = 1.2 × 10^−5^)(see Methods). Furthermore, we observed a similar trend using an independent set of plasma membrane proteins from the HPA database^23^ (see Methods; Supplementary Table 3). As the lack of addressing signal has been remarkably observed for dozens of cell surface moonlighting proteins^7^ performing alternate functions in other subcellular locations, we speculate that the formation of 3’UTR-protein complex could play a role in the unconventional plasma membrane translocation^11^ of the EMF proteins and more generally in the regulation of the localization of the EMF proteins.

**Table 2.** Numbers of nascent proteins in 3’UTR-protein complexes localized at the plasma membrane and without conventional translocation signals. Percentages denote the proteins retained compared to the previous column. Where indicated, Fisher’s exact tests were performed to test for statistical significance using as background the set of 7025 nascent proteins liable to be assessed for 3’UTR-protein complexes. The number of proteins in this background that are associated with the plasma membrane is 1877, of which 892 lack a membrane addressing signal. The Benjamini-Hochberg procedure was applied for multiple test corrections. Significance: ‘*’ indicates a P-value < 0.05; ‘**’ indicates a P-value < 0.01; ‘***’ indicates a P-value < 0.001.

| Protein group | Nascent in complex | of which, localized at the plasma membrane | of which, contain no membrane addressing signal |
| --- | --- | --- | --- |
| EMF | 128 | 45 (35%)* | 36 (80%) *** |
| Background | 7025 | 1877 (26.7%) | 892 (47.5%) |

To investigate the potential role of 3’UTR-protein complex formation in protein localization, we assessed whether EMF proteins found at the plasma membrane have been also observed in other ‘unexpected’ cellular locations. For this, we employed the PrOnto method^33^ which identifies dissimilar GO term pairs unlikely to occur among the annotations of the same protein, or among the annotations of interacting proteins (see Methods). Doing so, we discovered that more EMF proteins than expected are annotated to at least one pair of dissimilar Cellular Component GO terms (Fig. 5; 46.6% observed vs. 41.1% expected; empirical p-value = 0.022, 10,000 randomizations; Supplementary Table 6). Remarkably, the rarity of the observed proportion compared to expected (measured by the empirical P-value) increases when considering EMF proteins in 3’UTR-protein complexes (Fig. 5; 58.14% observed vs. 48.72% expected; empirical p-value = 9×10^−3^). This is particularly true for the subset of 36 EMF proteins found at the plasma membrane (Fig. 5; 92.3% observed vs. 60.07% expected; empirical p-value = 1×10^−4^). Indeed, this latter result postulates that 33 out of the 36 EMF proteins, besides being present at the plasma membrane, are also present at another ‘unexpected’ cellular location, demonstrating the great cellular localization versatility and fine-tune regulation of these proteins. Interestingly, 31 out of 33 of these proteins localize to the nucleus in addition to their plasma membrane localization, whereas only 5 of them have a nucleus addressing sequence (NES/NLS)^34^ (Supplementary Table 7).

**Figure 5.**
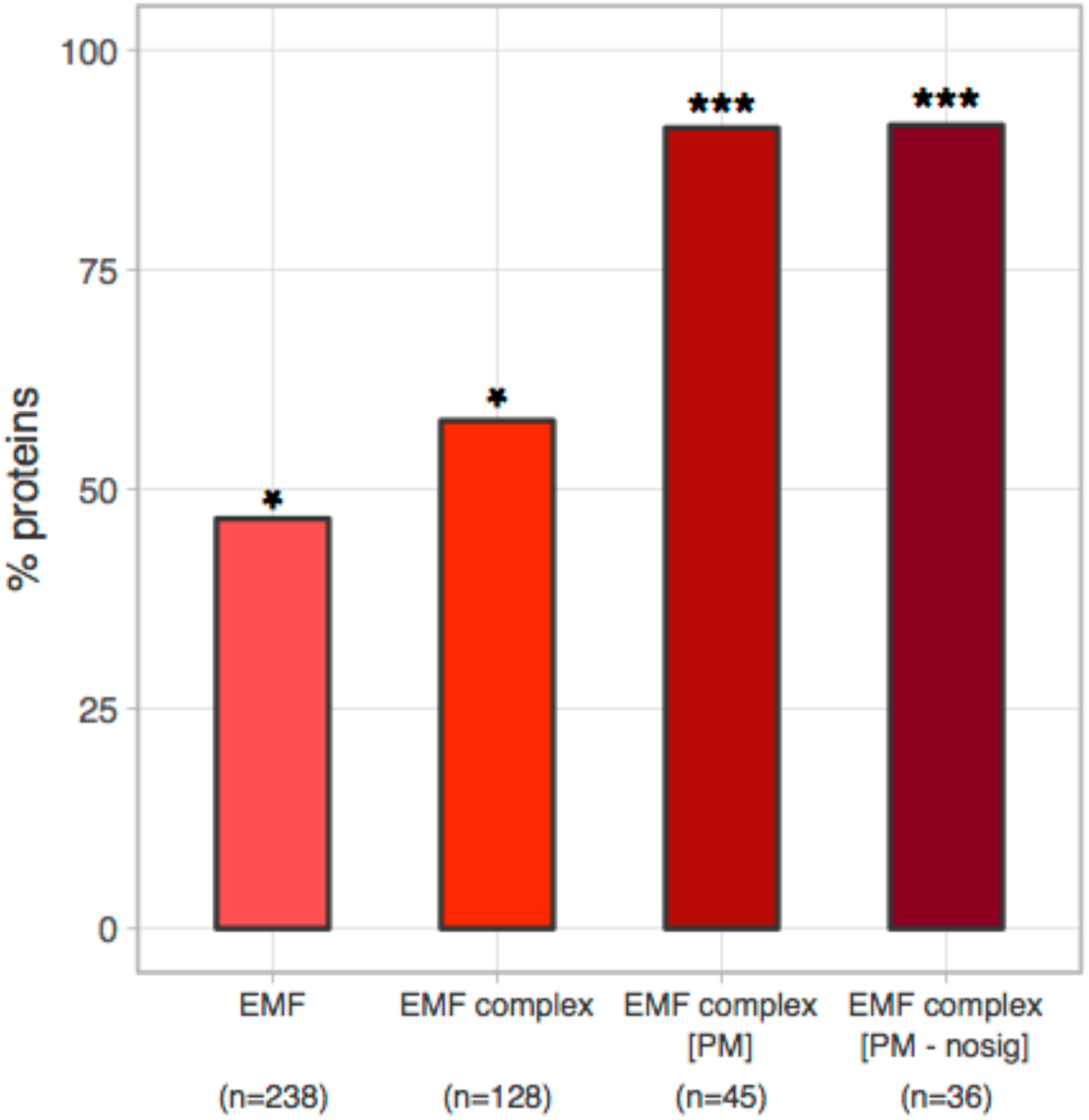
Percentage of proteins with GO term annotations to dissimilar cellular components. Dissimilar CC GO terms were determined with the PrOnto method. Each bar represents the percentage of proteins in the protein group annotated to at least one dissimilar GO term pair. We tested whether the observed proportion was higher than the expected values by sampling (10,000 times) sets of proteins from the human proteome of same size of the considered protein group and with identical distribution of annotated CC GO terms. ‘EMF’ refers to the 238 EMF proteins. ‘EMF complex’ represents the 128 EMF proteins in a 3’UTR-protein complex. ‘EMF complex [PM]’ refers to the subset of 45 EMF proteins which are found at the plasma membrane. ‘EMF complex [PM-nosig]’ refers to the subset of 436 EMF proteins which are found at the plasma membrane and lack addressing signal. Significance: ‘*’ indicates an empirical P-value < 0.05; ‘***’ indicates an empirical P-value < 0.001.

Overall, the vast majority of the EMF proteins found in 3’UTR-protein complexes and associated with the plasma membrane lack conventional membrane addressing signals and can also be found in the nucleus despite the absence of NES/NLS signals. This leads us to propose that 3’UTR-protein complexes could play a key role in the translocation of these extreme multifunctional proteins, not only by participating in protein transport to the plasma membrane as described for CD47^11^ but more generally, in protein trafficking between different subcellular compartments.

### 3’UTR-protein complexes mediate multifunctional protein trafficking linked to signaling and cell migration processes

If the 3’UTR-protein complexes participate in protein transport between subcellular compartments, EMF protein functional annotations should reveal and highlight the biological processes requiring their formation. For this, we performed a GO term analysis on the 128 EMF proteins contained in the 1411 3’UTR-protein complexes. Strikingly, the significant enrichments in membrane-related cellular components (“adherens junction”, “extrinsic component of membrane”, “extracellular exosome”, “whole membrane”), along with “nucleoplasm” and “cytosol” evoke cellular trafficking (Fig. 6a). In addition, the significant enrichments in functions related to “phosphatidylinositol phosphorylation”, “kinase binding”, “regulation of calcium ion transport into cytosol” and “calcium-mediated signaling” point towards signaling processes. Further GO term analysis on the intermediate proteins linking the EMF proteins to the RBPs within the 3’UTR-complexes, confirm the significant enrichments of trafficking- and signalling-related terms such as “regulation of clathrin-dependent endocytosis”, “endocytic vesicle membrane”, “receptor internalization”, “ephrin receptor-” and “steroid hormone receptor-binding activities”, and “protein kinase activity” (Fig. 6b). Finally, 3’UTR-complexes could participate to neuronal processes, as suggested by enriched terms such as “synapse”, “learning or memory” and “dendrites” among EMF and intermediates proteins, respectively. On the other hand, vascular and muscular processes through cell migration are also suggested by enriched terms like “cellular response to angiotensin”, “regulation of blood vessel endothelial cell migration”, “positive regulation of smooth muscle cell migration”, “muscle hypertrophy”, “ventricular septum development”, and “homeostasis of number of cells” among intermediate proteins. At the molecular function level, it is important to note that when EMF proteins are enriched in “kinase binding” proteins, the intermediate proteins with which they interact are expectedly enriched in “protein kinase activity”, “non-membrane spanning protein tyrosine kinase activity” and “protein kinase regulator activity”, suggesting that the recruitment of kinases as intermediate proteins is important for EMF nascent protein regulation or fate. Interestingly, conversely to EMF proteins, intermediate proteins are enriched in proteins containing potential NES/NLS signals according to NLSdb^34^ (OR=1.6, two-sided Fisher’s Exact Test, P-value = 3.70 × 10^−5^) and in the GO term “nuclear localization sequence binding” (P-value= 1.1×10^−2^, Fig6b), suggesting their active role in translocating EMF proteins. Furthermore, RBPs are enriched in “mRNA-containing ribonucleoprotein complex export from nucleus” GO term (P-value=5.3×10^−11^, see Supplementary Table 7), confirming the implication of the 3’UTR-complexes in protein trafficking between cellular compartments, more particularly across the nuclear membrane.

**Figure 6.**
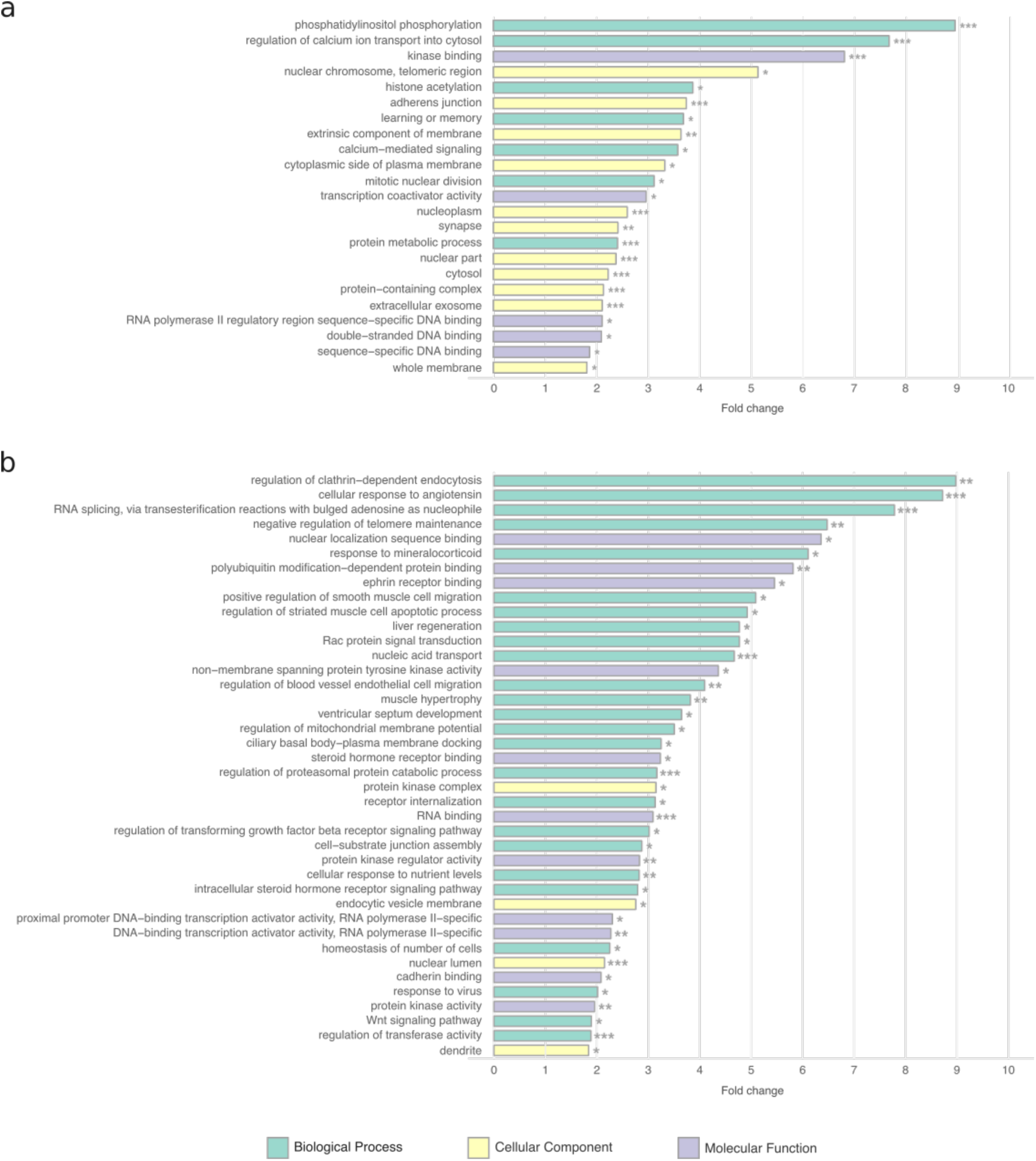
Functional enrichment analysis of EMF and intermediate proteins. Over-represented GO terms among **(a)** EMF nascent proteins and **(b)** their intermediates. Functional enrichment analysis was performed using the g:Profiler R package (see Methods). Proteins in the human PPI network were used as statistical background. Significance: ‘*’ indicates a FDR < 0.05; ‘**’ indicates a FDR < 0.01; ‘***’ indicates a FDR < 0.001.

### Predicted 3’UTR-protein complexes could explain protein multifunctionality

If 3’UTR-complexes formation contributes to protein multifunctionality, distinct 3’UTR-complexes assembled with several RBPs and different intermediates proteins, could provide the molecular environment for the EMF protein to perform different functions by interacting with different protein partners. In order to test this hypothesis, our reasoning was the following.

The EMF proteins used in this study were identified at the intersection of at least two network modules^35^ involved in dissimilar cellular processes, according to GO Biological Process annotations^17^ (Fig 7a). If the formation of a 3’UTR-protein complex contributes to the multifunctionality of a given EMF protein as recently shown for BIRC3^16^, we expect that the following conditions are met: *(i)* the EMF protein participates to different 3’UTR complexes (i.e. at least two); *(ii)* the recruited protein partners, i.e. the different intermediate proteins, found in each of these complexes should belong to different network modules involved in dissimilar biological processes (Fig 7a). We found that 82% of the EMF proteins present in 3’UTR complexes fulfill the first condition (i.e. 105 out of 128) and, among these, 44% (46 out of 105) meet the second one (see Methods), thus proposing a molecular scenario for the multifunctionality of almost half of the EMFs involved in 3’UTR-protein complexes.. Similarly, since a change of function is often associated with a change in cellular localization, we checked how many of the 105 EMF proteins present in two or more 3’UTR complexes, have at least two intermediate proteins belonging to different network modules annotated with dissimilar GO cellular localizations (see Methods, Fig. 7a). This was the case for 59% of these EMF proteins (i.e. 62 out of 105) and remarkably, 37 of the 46 EMF proteins with intermediate proteins belonging to network modules with dissimilar Biological Process annotations (i.e., 80,4%, Fig. 7b). Overall, the formation of 3’UTR-protein complexes seems to provide two important prerequisites for multifunctionality: diversity in interaction partners and distinct subcellular locations.

**Figure 7.**
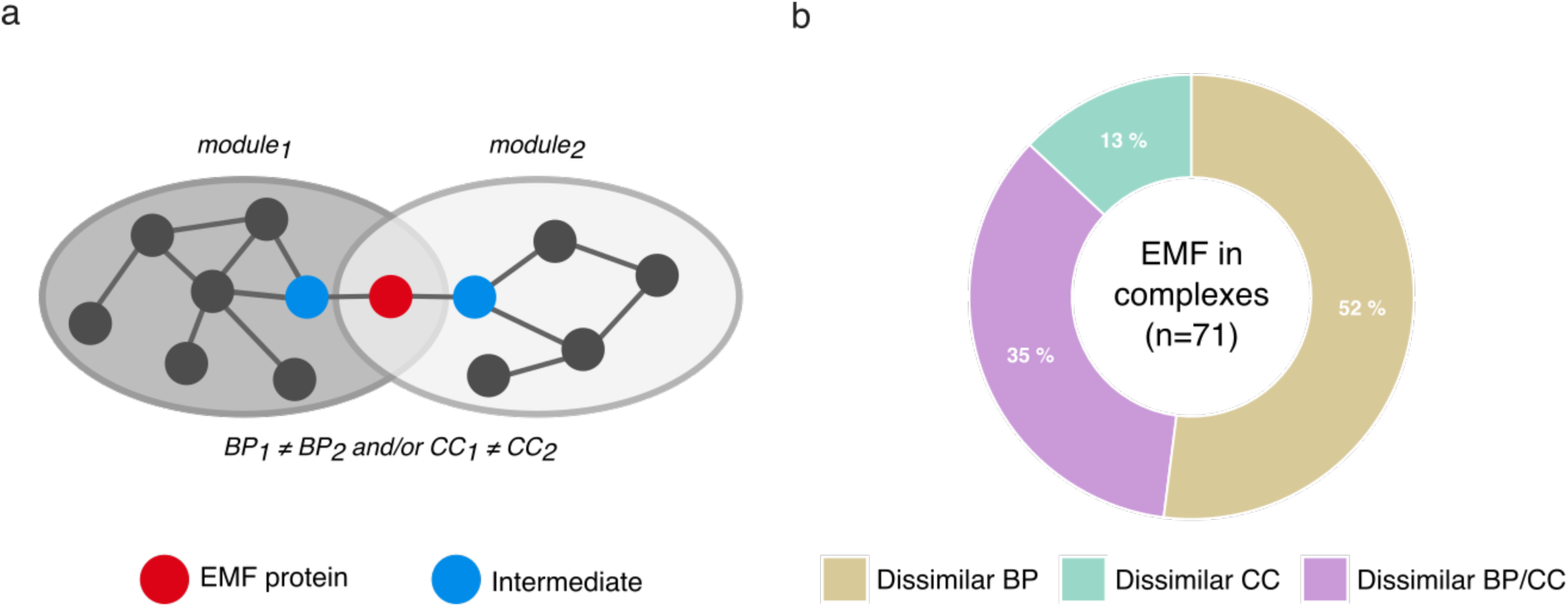
Network module annotations support a 3’UTR-complex role in multifunctionality. **(a)** Network modules containing EMF proteins were annotated with GO biological process (BP) and cellular component (CC) terms. In a pairwise fashion, for each EMF protein, we check if any of its intermediate proteins were present in distinct network modules with dissimilar BP (shades of green) or CC (shades of yellow) annotations. **(b)** The majority of EMF with intermediate proteins belonging to network modules with dissimilar BP annotations also have intermediate proteins present in network modules with dissimilar CC annotations.

### Archetypal moonlighting proteins are predicted to form 3’UTR-protein complexes

As our analysis has been performed on a dataset of candidate proteins to extreme multifunctionality, we aimed at confirming that experimentally verified moonlighting proteins could participate in 3’UTR-protein complexes. For this, we first verified that proteins of the reference dataset available in the MoonDB^22^ and the MoonProt^36^ databases (47 and 65 literature-curated moonlighting proteins, respectively), form such complexes. Indeed, up to 52% of the known moonlighting proteins do so, a proportion that is similar to the one obtained for the 238 predicted EMF proteins (54%, Table 1). Moreover, the number of predicted complexes is significantly higher than expected by chance for both moonlighting proteins datasets (Supplementary Table 4).

Interestingly, our set of EMF proteins in 3’UTR-protein complexes includes the alpha-enolase (ENO1), a well-known cytosolic glycolysis enzyme that moonlights at the cell surface as a receptor and activator of plasminogen, being thereby involved in cell migration, tissue remodeling, apoptosis and tumorigenesis^37^. Notably, ENO1 lacks a signal sequence and the mechanism leading to its translocation between subcellular compartments is yet unknown^37^. We found that ENO1 forms 8 different 3’UTR-protein complexes with different RBPs and intermediate proteins with different tissue expression.

Interestingly, one of them is only predicted in muscles (heart, skeletal and smooth muscles). It contains DDX6, an RNA helicase involved in muscular dystrophy^38^ as RBP and Desmin (DES), forming myofibrils and linking them to the cytoskeleton, the nucleus, mitochondria, and the plasma membrane, as intermediate. It is thus tempting to speculate that this complex participates or mediates the plasminogen receptor function of ENO1 in muscle regeneration and muscle injury recovery^37^ by localizing it at the muscle cell surface. Finally, the fact that 4 out of 5 RBPs involved in the 3’UTR-ENO1 complexes bind only the longer ENO1 3’UTRs, as described for CD47^11^ and BIRC3^16^ and that 4 out of 7 intermediates proteins are annotated to both intracellular and extracellular locations, increases our confidence in the predictions.

Likewise, we also predicted a 3’UTR-protein complex formed with the nascent protein RHAMM/HMMR, a regulator of the stability of the mitotic spindle in normal cells^8, 10^ that acts as an extracellular CD44-ligand promoting cell motility and invasion in cancer cells^10^, IGF2BP2, involved in mRNA storage and transport, and the Dynactin subunit 1 (DCTN1) as intermediate, a protein present at the “spindle” as well as at the “membrane” and notably involved in the transport of organelles and vesicles by tethering the dynein cargo to the microtubule^39^. We can thus hypothesize that the 3’UTR-RHAMM complex plays a role in the molecular mechanism allowing the extracellular localization of RHAMM in tumor cells.

Finally, in addition to enolase, almost all other glycolytic enzymes have been shown to display moonlighting activities unrelated to glycolysis. These additional functions often explain the phenotypes observed in the metabolic disorders caused by the dysregulation of these enzymes^40^. Very interestingly, we found that not only proteins involved in the “glycolytic process” are enriched among the nascent proteins of the interactome present in the 3’UTR-protein complexes (P-value= 3.0×10^−3^), but that all the enzymes of the glycolysis pathway are involved in 3’UTR-protein complexes, further pushing the idea that 3’UTR-protein complex formation plays a role in protein multifunctionality.

### The formation of 3’UTR-protein complexes is a prevalent mechanism at the interactome-scale

What is the prevalence of the 3’UTR-protein complexes formation? Is it a widespread mechanism in the cell and not limited to known moonlighting and EMF proteins? As shown in Table 1, our predictions revealed that 17% of the proteins of the interactome have the potential to be involved in at least one 3’UTR-protein complex, i.e., ∼3 times more than expected randomly (empirical P-value=0.091). Interestingly, we found that nascent proteins involved in 3’UTR-protein complexes show many of the features previously described for the EMF proteins: they are enriched in translocating and multilocalizing proteins, although largely devoid/depleted in membrane addressing features and signal peptides. Additionally, the enriched GO annotations among the 2354 nascent proteins and their intermediate proteins recapitulate the biological processes and the subcellular compartments found as enriched for EMF proteins and their intermediates including ion homeostasis, blood vessel endothelial cell migration, cell-cell signaling, extrinsic and cytoplasmic membrane parts, coated membrane, extracellular exosome and as well as nuclear part. However, proteins localized within the membrane and annotated “intrinsic component of membrane” and “integral component of plasma membrane” are intriguingly underrepresented among the nascent proteins of the 3’UTR-protein complexes. This result may either reflect a methodological bias — as transmembrane proteins are under-represented in two-hybrid screen results, which contribute significantly in terms of interactions in our human PPI network — or the fact that as 3’UTR-protein complexes are involved in protein trafficking, they leave aside integral proteins of the membrane.

Overall, our results suggest that the formation of 3’UTR-protein complexes could be a prevalent mechanism in the cell that could promote protein multifunctionality through protein trafficking and/or subcellular localization change for almost 20% of the proteins of the interactome.

## Discussion

What are the molecular mechanisms that enable the functional changes of moonlighting proteins? Could the formation of 3’UTR-protein complexes explain the unexpected membrane localization of some multifunctional/moonlighting proteins in the absence of membrane addressing signals? Could these ribonucleoprotein complexes promote the multifunctionality of some proteins by recruiting different interaction partners that are necessary to their switch of function?

The regulation of protein localization and function by the formation of ribonucleoprotein complexes mediated by 3’UTRs, have been proposed for two proteins, namely CD47^11^ and BIRC3^16^. However, the prevalence of the formation of such complexes is still unknown. In order to circumvent the current scarcity of 3’UTR-complexes that hinders any large-scale analysis, we propose a computational mapping strategy that allows inferring, for the first time, those ribonucleoprotein complexes. For this, we took advantage of the availability and the combination of two network data types — 3’UTR-protein and PPI networks —, covering a large part of the human transcriptome and proteome. Our inferences allowed us to both provide a detailed analysis of the characteristics of the 3’UTR-protein complexes as a group and evaluate the prevalence of their formation at the interactome level. We indeed discovered that as much as 54% of the extreme multifunctional proteins and up to ∼20% of the whole interactome proteins could be involved in such complexes, thereby underlining their potential high extent and cellular importance. However, our inferences are notably sensitive to the comprehensiveness of the analyzed data since cellular 3’UTR-protein complexes that are formed by molecular interactions not present in the analyzed datasets cannot be found. Indeed, while experimental human PPI networks cover most interacting proteins^41^, public interaction databases may not include all the interactions known in the literature or recently discovered. In addition, when building the PPI network to be analyzed, some interactions may have been filtered to reduce false-positive predictions as much as possible. Indeed, our large-scale PPI network contains only direct binary interactions, while indirect interactions were not considered (e.g., those identified by co-immunoprecipitation techniques). This explains why the ELAVL1-SET-CD47 and the different BIRC3 protein complexes have not been found by our approach. In addition, taking into account that current RBP-3’UTR interaction datasets contain data for only a subset (less than 400 RBPs^28^) of an increasingly growing number of proteins is interacting with RNAs (which may amount to as many as 2000 RBPs^42, 43^), we may have underestimated the number of existing 3’UTR-protein complexes in human cells. In addition, the contribution of ribonucleoprotein complexes to a number of cellular processes may have been largely overlooked so far, due to the fact that usual experimental methods to identify cellular macromolecular complexes routinely use an RNA nuclease step before protein purification, thereby hindering the possible detection of RNA components in protein complexes^44^. Computational approaches can help to overcome this drawback, as suggested by our results. Nevertheless, this study provides for the first time an extensive overview of the 3’UTR-protein complex formation for a subset of human proteins, predicting that a sizeable amount of such cellular complexes can be formed, employing a large variety of RBPs and intermediate protein components.

The diversity in subcellular localization and protein partners are two main protein multifunctionality determinants possibly influenced and driven by 3’UTRs. 3’UTRs are generally described as responsible for the localization of their cognate mRNAs in asymmetrical and polarized cells such as neurons, where localized translation modifies the nearby proteome in response to external cues^12, 45^. This is mediated by RBPs that interact with motor proteins, allowing the transport of mRNA along actin cables. On the other hand, 3’UTRs also mediate PPIs, as in the case of the 3’UTR-protein complexes described in our study. Notably, the enrichments in the GO terms “nucleic acid transportation”, “synapse”, “dendrite”, coupled to the function “learning and memory” among EMF and intermediate proteins suggest that these two 3’UTR functions, mRNA transport and protein complex scaffolding, are related and intermingled.

What could be the roles of the 3’UTR-protein complexes in protein localization change? In the CD47 case, protein localization to the plasma membrane depends on the intermediate protein SET and its protein partner, active RAC1^11^. Very interestingly, on a global scale, the enrichment in “Rac protein signal transduction” annotation among intermediate proteins that we observe could extend this dependency to a subset of predicted 3’UTR-protein complexes containing EMF proteins. This would suggest that small GTPases signaling generally contribute to the protein transport mediated by the formation of the 3’UTR-complexes. In addition, given the role of RAC signaling in actin remodeling and cell migration^46^, it is interesting to note that other proteins involved in the regulation of cell migration are also enriched among intermediate proteins (Fig. 6b). This underlines a possible involvement of the trafficking mechanism we describe in cell migration processes.

Most multifunctional proteins predicted to be translocated by the 3’UTR-protein complexes do not contain the corresponding conventional addressing signals in their sequence. However, most of the multifunctional proteins participating in 3‘UTR-protein complexes are located in the nucleus and at the membrane. The 3’UTR-complex and more particularly, the intermediate protein could help to address properly the nascent protein. Indeed, the fact that intermediate proteins are enriched in NLS/NES signals suggest that they may contribute to the nuclear localization of their protein partners without NLS, which would then behave as cargos. Moreover, according to GO term enrichment analysis, some intermediate proteins bind NLS sequences. These are importins and transportins able to bind the nuclear pore complex and involved in nuclear protein import and export. Interestingly, these intermediates could contribute to the nuclear localization of their protein partners devoid of NLS, as the importance of the three-dimensional context rather than just the presence of the NLS has been recently illustrated for the recognition of cargos by importins^47^. Together with the fact that intermediates are also enriched for endocytosis, receptor binding, and internalization, these results confirm the role of the 3’UTR-protein complexes in protein transport between the subcellular compartments at the cell scale.

Protein multifunctionality can be controlled by the formation of 3’UTR-protein complexes. Depending on the cell type and cellular conditions, alternative 3’UTR isoforms may bind different RBPs, that may interact with different intermediates, ultimately leading to different 3’UTR-protein complexes containing the same nascent protein but serving different functions. We showed this is the case for half of the EMF proteins contained in 3’UTR-complexes. Notably, it also has been shown for BIRC3, originally known as an E3 ubiquitin ligase, but which has been recently discovered to be involved in protein trafficking, chromatin regulation, and mitochondrial processes as well, through the 3‘UTR-protein complexes it belongs to^16^.

The assembly of 3’UTR-complexes may provide the molecular context and proximity necessary for protein moonlighting functions. Indeed, a large number of intermediate proteins are kinases that can modify nascent proteins, illustrating the fact that moonlighting functions may be partly commanded by short linear motifs^17, 48^. Recently, it has been shown that 4 out of 9 *S. cerevisiae* protein complexes that form co-translationally^49^, possibly according to a 3’UTR-complex formation model^50^, involve 6 known multifunctional and moonlighting yeast proteins^49^. Interestingly, 2 of those, MetRS and GluRS, possess nuclear and mitochondrial localization signals, which are only revealed when they proteins are not in complex with each other^51^, indicating that their co-translational complex assembly – and possibly 3’UTR-protein complex formation – may regulate their moonlighting function.

Overall, our data bring the first large-scale prediction and analysis of 3’UTR-protein complexes. We propose that the formation of 3’UTR-protein complexes is a widespread phenomenon allowing *(i)* the transport of proteins between subcellular compartments in the absence of conventional addressing signals, and *(ii)* the assembly of multiple complexes that sustain the multifunctionality of proteins, thus representing a plausible molecular mechanism for the regulation of protein moonlighting functions.

## Methods

### Protein-protein interaction network, EMF proteins and protein groups

Predicted human extreme multifunctional (EMF) proteins (238 proteins) and a human PPI network (14046 proteins, 92348 interactions) were downloaded from MoonDB 2.0^22^ (http://moondb.hb.univ-amu.fr/). The human PPI network was constructed by interactions retrieved from the PSICQUIC web service^52^ on January 2018, as described in^17^, and does not contain ‘self-interactions’. Network modules were extracted from the PPI network using OCG^35^, a clustering algorithm that allows proteins to belong to more than one cluster. These network modules are available on MoonDB 2.0. Briefly, EMF proteins (238 proteins) are proteins that belong to two or more network modules whose Gene Ontology (GO) term annotations (‘Biological Process’) contain at least two terms that are dissimilar to each other according to PrOnto^33^. GO term annotations and ontologies were collected from the Gene Ontology Consortium^53^ on December 2017. Analysis involving the ‘proteome’ protein group used a human proteome (20349 proteins) retrieved from UniProt (‘reviewed’ proteins only) on June 2018^54^.

### Datasets of 3’UTRs and polyadenylation sites

Ensembl v90 spliced 3’UTR sequences for all human transcripts were downloaded from the Ensembl BioMart service^25^. The maximum 3’UTR length was calculated for each protein in the human proteome (UniProt AC) by selecting the longest 3’UTR among all transcripts encoding for a certain protein. Genome-wide polyadenylation sites for human were downloaded from APADB v2^26^ as well as PolyASite version r1.0^27^, on December 2017. APADB polyadenylation sites per kb were calculated for proteins produced from transcripts with 3’UTRs longer than 1000 nt, taking into account the length of the longest 3’UTR. For PolyASite, polyadenylation sites on the terminal-exon “TE” category were considered. Gene names and Ensembl transcript IDs were converted to UniProt AC using the UniProt ID mapping tool^54^.

### RBP-3’UTR interaction network

Interactions between RBPs and 3’UTRs were retrieved from the Atlas of UTR Regulatory Activity (AURA) v2.4.3 database (AURAlight dataset) on January 2018^28^. The AURA database contains interactions between 3’UTRs and RBPs collected and mapped from various experiments, including several types of cross-linking and immunoprecipitation (CLIP) methods. Gene and coding-transcript identifiers were mapped to reviewed UniProt ACs using UniProt ID cross-referencing files (HUMAN_9606_idmapping.dat)^54^. Only interactions involving proteins present in the PPI network were used.

### Prediction of 3’UTR-protein complexes

3’UTR-protein complexes were predicted with in-house Python v2.7 scripts using the protein-protein and RBP-3’UTR interaction networks described above.. Each 3’UTR-protein complex includes: *(i)* an interaction between the RBP and the 3’UTR, *(ii)* an interaction between the intermediate protein (i.e. the protein which interacts with both the RBP and the nascent protein) and the nascent protein, *(iii)* an interaction between the intermediate protein and the RBP. We only considered the presence of one intermediate protein, and complexes formed without any intermediate protein were not examined (*i.e*., RBP interacting directly with the nascent protein). Since the PPI network used does not contain self-interactions, the intermediate protein must be different from the nascent and the RBP. 3’UTR-protein complexes were detected for EMF as well as all proteins in the human PPI network. Only proteins with (*i)* PPIs, (*ii)* 3’UTR-RBP interactions and (*iii)* presence in at least one HPA tissue (see below) were liable to be assessed for 3’UTR-protein complexes as ‘nascent’ proteins. 3’UTR-protein complexes were further filtered according to protein tissue presence, as described below.

### Protein tissue presence filter

Tissue protein presence from Human Protein Atlas (HPA) version 18 (January 2018)^29^ was used to filter 3’UTR-protein complexes. This dataset contains data on 58 normal tissues. Information on cell type associated with tissue names was not used in this study. Proteins with reliability score (level of reliability of the protein expression pattern) indicated as ‘uncertain’ and proteins with presence level ‘not detected’ were excluded. We only considered 3’UTR-protein complexes where all proteins of the complex are present in at least one of the 58 tissues. Gene names and Ensembl Gene IDs were converted to 13044 reviewed UniProt AC using the UniProt ID mapping tool^54^.

### 3’UTR usage in predicted complexes

Sequences and hg19 genomic coordinates of 3’UTRs interacting with RBPs were taken from the AURA database (file: UTR_hg19.fasta, 65,285 sequences) on December 2018. As AURA does not provide cross-references to Ensembl identifiers, we used BEDTools^55^ intersect with parameters “*-s -r -f 1.0*” (to ensure strendness and a perfect match) to map AURA UTR coordinates on GENCODE release 19 (GRCh37.p13) gene annotation file. In doing so, we were able to associate 62,898 UTR sequences (96% of the total) to the corresponding protein-coding genes. UTR sequence redundancy was reduced at 100% identity using the CD-HIT algorithm^56^.

### Proteins localized in the plasma membrane

Plasma membrane proteins were retrieved from two datasets: (*i)* UniProt^54^, querying reviewed *Homo sapiens* proteins with the GO term ‘plasma membrane’ (GO:0005886) annotated by ‘any manual assertion’ method (4602 proteins) and (*ii)* plasma membrane proteins experimentally detected by HPA version 18^23^, querying for the subcellular locations ‘plasma membrane’ and ‘cell junctions’ (1734 genes mapped to 1776 UniProt ACs using the UniProt ID mapping tool). Note that both datasets include proteins that are integral to the plasma membrane (e.g. receptors) as well as peripheral membrane proteins that may attach to integral membrane proteins or penetrate the peripheral regions of the membrane (e.g., receptor-interacting proteins). Information on the presence or absence of signal peptide, lipidation sites, and transmembrane domains was obtained from UniProt on January 2019^54^. The set of 7025 nascent proteins liable to form 3’UTR-protein complexes (i.e. having protein-protein and protein-RNA interactions, as well as present in HPA), even though not enriched in plasma membrane proteins, contain a higher proportion of proteins localized in plasma membrane without a signal peptide or transmembrane domains than the proteome (52.4% versus 31.4%, using UniProt data). Thus, to avoid potential biases, statistical comparisons were done against this set of proteins instead of the proteome in plasma membrane-related analysis.

### Dissimilar cellular component GO term analysis

The PrOnto method^33^ was used to identify ‘cellular component’ (CC) GO term pairs that are unlikely to occur in the same protein (association probability) or in interacting proteins (interaction probability). PrOnto probabilities were recalculated for CC GO term annotations collected from the Gene Ontology Consortium^53^ on December 2017, including terms ‘Inferred from Electronic Annotation’ (IEA). GO term pairs with PrOnto association and interaction probabilities <0.05 were considered dissimilar. EMF protein dissimilar GO terms can be consulted on the related MoonDB v2.0 database. We tested whether certain protein groups have more proteins with dissimilar terms than expected. Since the ability to find dissimilar GO term pairs depends on the number of GO terms annotated to a protein, we control this by sampling (10,000 times) sets of proteins from the human proteome of the same size of the protein group in question and with the same distribution of numbers of CC GO terms annotated.

### Gene Ontology enrichment analysis

The Gene Ontology (GO) term enrichment analysis was performed using the g:Profiler R package^57^. We only considered enrichments where the minimum query/term intersection size was =>5, and where the P-value was <0.05 after correction using the Benjamini-Hochberg procedure. In all g:Profiler analysis, electronic GO annotations (i.e. with the evidence code IEA) were considered.

### Modules with dissimilar annotations

We retrieved from MoonDB v2.0 the list of network modules and the pairs of *(i)* dissimilar functions (i.e. GO Biological Process terms) annotating the modules used for the identification of the EMF proteins, and *(ii)* dissimilar GO Cellular Component annotating the modules. We selected EMF proteins present in at least two predicted 3’UTR-protein complexes. Subsequently, for each selected EMF protein, we checked whether any pair of intermediate proteins were present in distinct network modules, annotated to dissimilar GO BP, or dissimilar GO CC, or both.

## Acknowledgments

We would like to thank Lionel Spinelli (TAGC) for critically reading the manuscript and Philippe Pierre (CIML) for fruitful scientific discussions. The project leading to this publication has received funding from the Excellence Initiative of Aix-Marseille University - A*MIDEX, a French “Investissements d’Avenir” program (to CB).

## Author Contributions

AZ and AT performed preliminary analyses. DR and AT developed the analysis pipelines. DR and AZ gathered the final data. DR, AP, and AZ performed the experiments. AZ and CB conceived the study, with inputs from DR and LS along the project. DR, AZ, and CB wrote the manuscript, helped by the other authors. AZ and CB supervised the work. CB secured funding. All authors approved the final draft.

## Supplementary Figures

**Figure S1.**
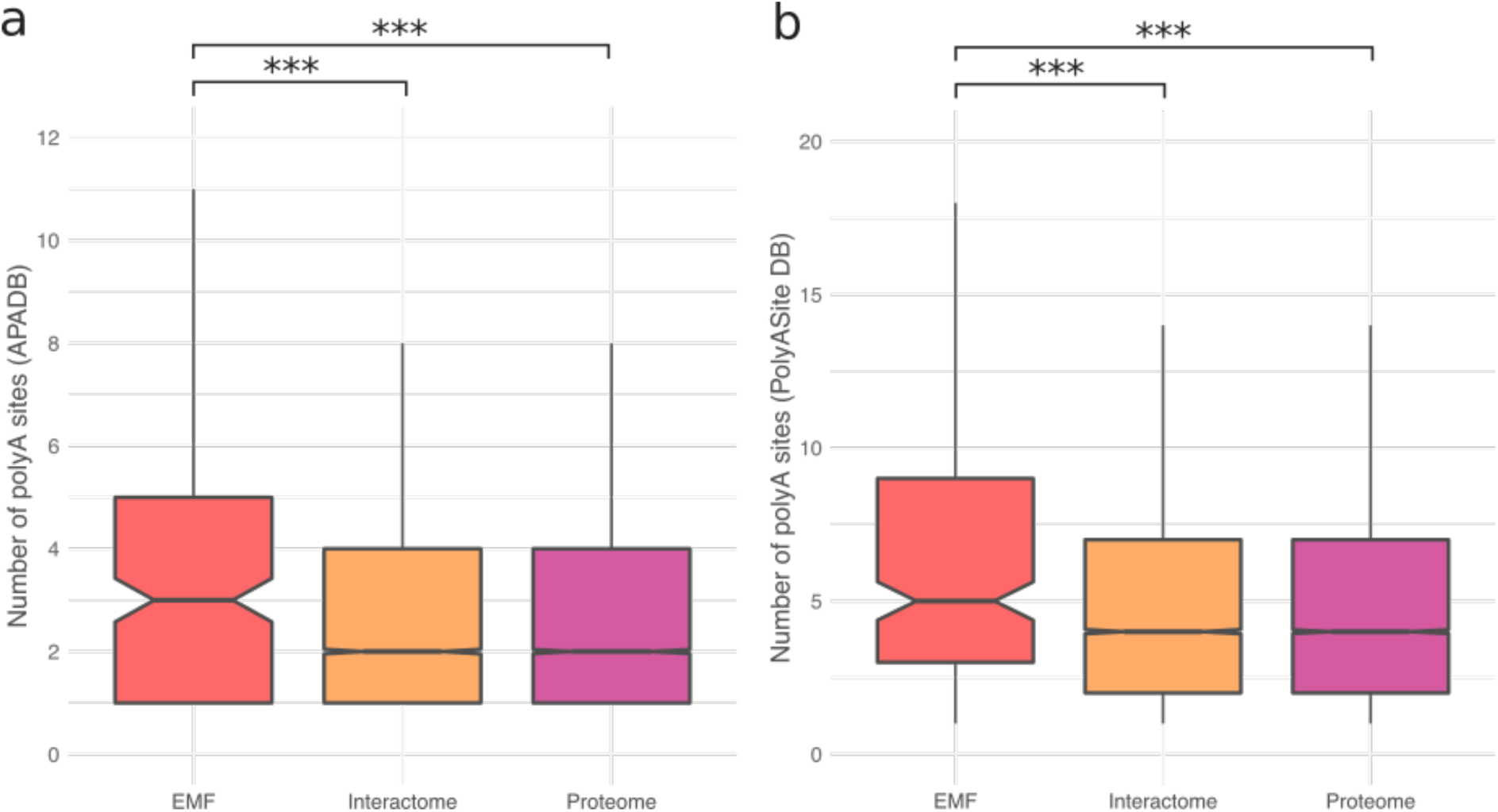
Alternative polyadenylation (APA) sites in EMF 3’UTRs. Comparison of the number of APA sites in the 3’UTRs of mRNA encoding EMF proteins and those encoding the other groups of proteins. **(a)** APA sites from APADB, Kruskal-Wallis rank sum test, P-value = 6 x 10^−8^. **(b)** APA sites from the PolyASite database, Kruskal-Wallis rank sum test, P-value < 2 x 10^−16^. Mann-Whitney U tests were performed to test for pairwise statistical significance. The Benjamini-Hochberg procedure was applied for multiple test correction. Significance: ‘***’ indicates a FDR < 0.001.

**Figure S2.**
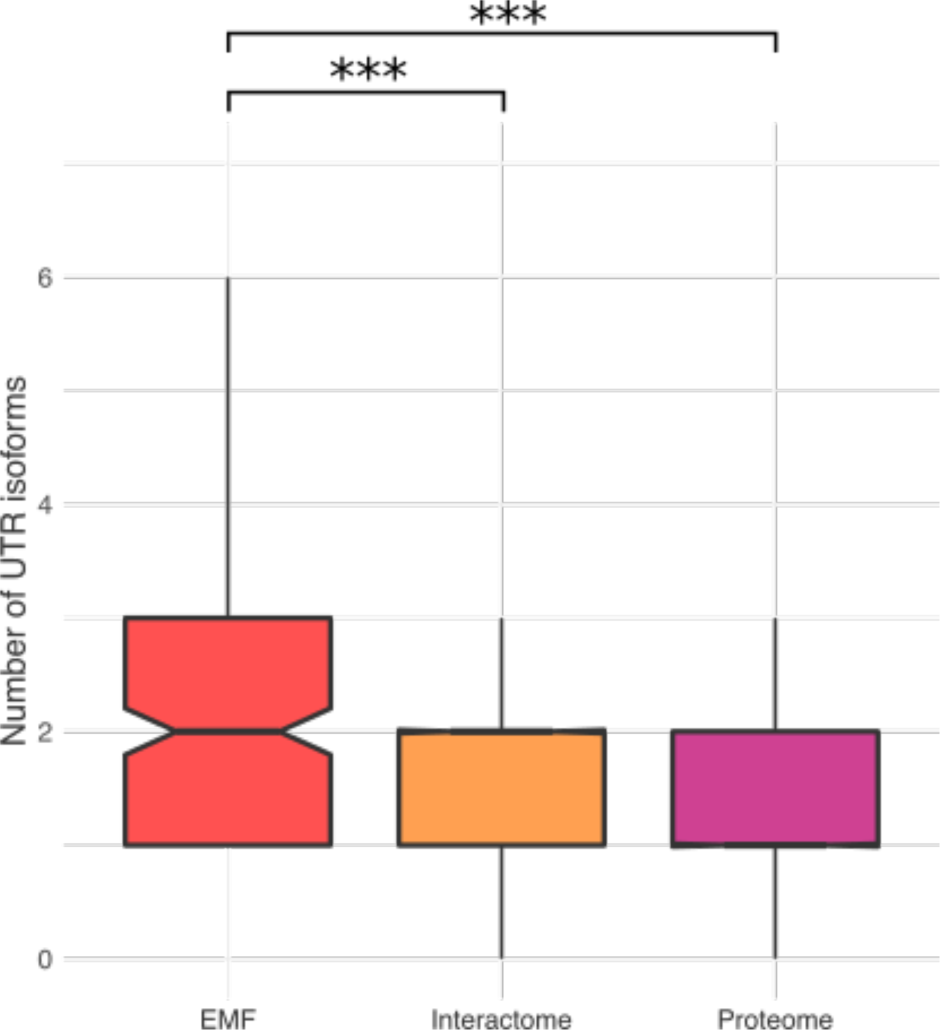
3’UTR lengths. Comparison of the lengths the of 3’UTRs of mRNA encoding EMF proteins and those encoding the other groups of proteins. Kruskal-Wallis rank sum test, P-value < 2 x 10^−16^. Mann-Whitney U tests were performed to test for pairwise statistical significance. The Benjamini-Hochberg procedure was applied for multiple test correction. Significance: ‘***’ indicates a FDR < 0.001.

**Figure S3.**
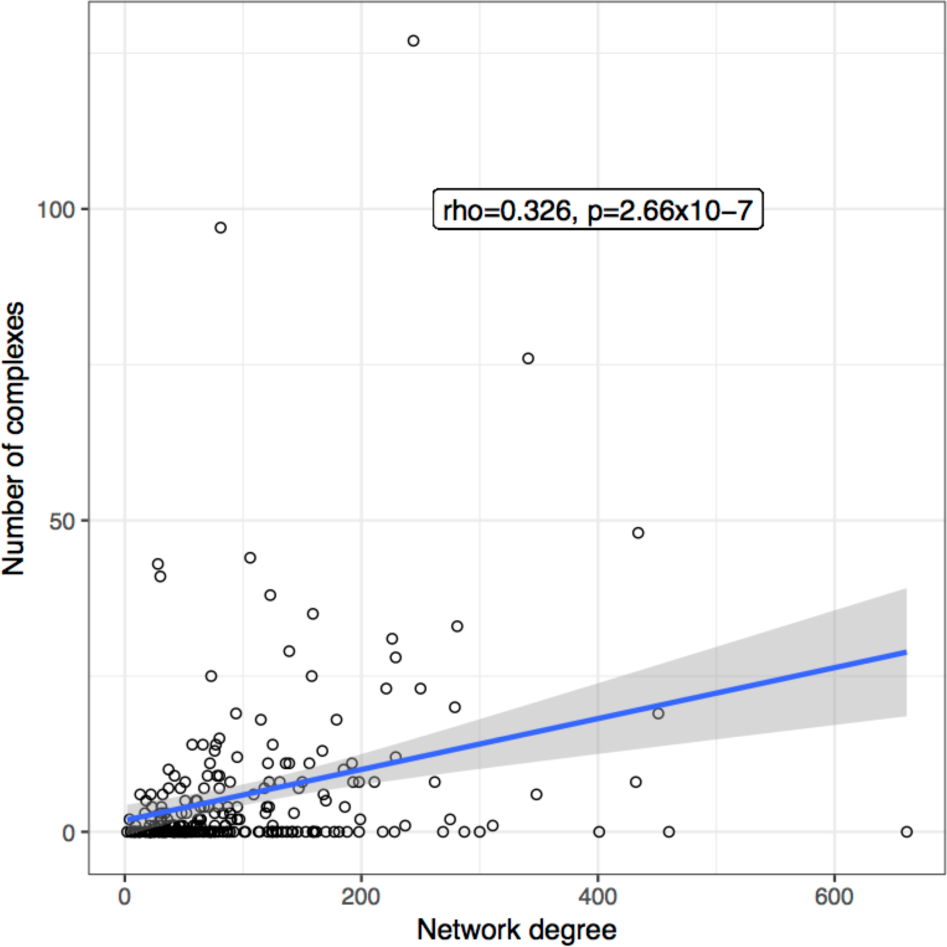
Correlation between EMF network degree and number of complexes. The network degree of EMF proteins represents the number of interaction partners in the human binary interactome. The correlation with the number of complexes in which EMF proteins participate was computed using the Spearman’s rho coefficient. The blue line represents a linear regression fit and the gray area is its 0.95 confidence interval.

**Figure S4.**
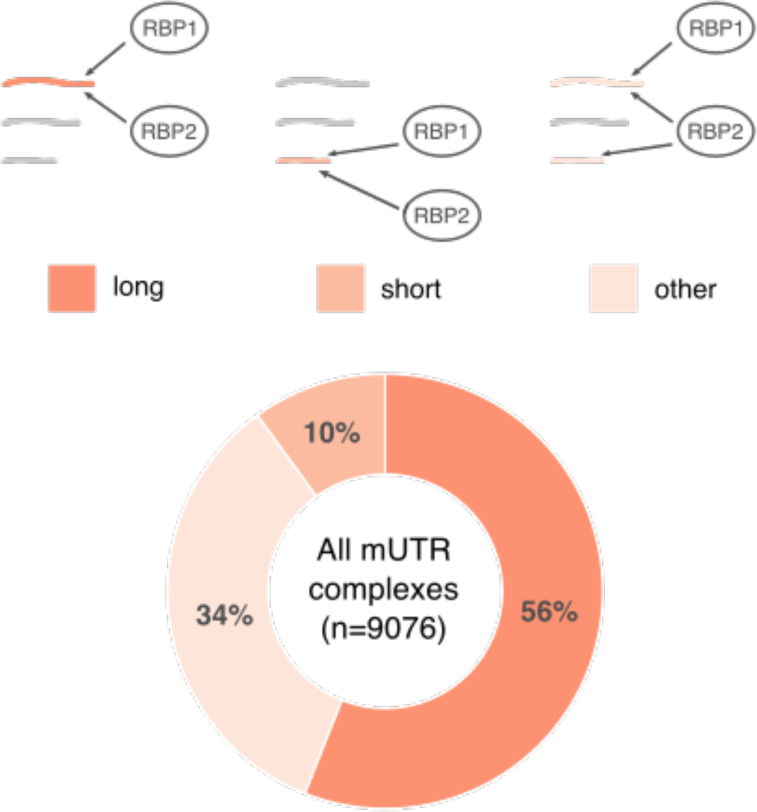
Preferential isoform usage in mUTR predicted complexes in the binary interactome. Nascent mUTR proteins were classified based on the length category to which their RBP-binding 3’UTR(s) belong to the longest or the longer isoforms are bound to RBPs (i.e, “long” category); the shortest or shorter isoforms are bound to RBPs (i.e., “short” category); RBPs bind all isoforms, or a substantial fraction of them, without a clear preference for long or short isoforms (i.e., “other” category).

## Supplementary Tables

**Supplementary Table 2.** Number of 3’UTR-protein complexes expected by chance. The total number of 3’UTR-protein complexes expected by chance was measured on protein-protein interaction networks that had their protein labels randomly shuffled without replacement (e.g. ‘protein 1’ becomes ‘protein 2’, ‘protein 2’ becomes ‘protein 55’ etc.). Empirical p-values were calculated for the hypothesis that the number of 3’UTR-protein complexes formed using the real protein-protein interaction network is higher than the null distribution of 10,000 randomizations. This analysis was performed for the whole set of possible nascent proteins (7088 proteins). The mean number of total complexes and its standard deviation (in parenthesis) over the randomizations is presented.

| Protein group | 3'UTR-protein complexes | 3'UTR-protein complexes in reshuffled networks | Empirical p-value |
| --- | --- | --- | --- |
| EMF | 1411 | 117 (67.2) | 0.0001 |
| Interactome | 9657 | 3478.9 (979.4) | 0.091 |

**Supplementary Table 3.**
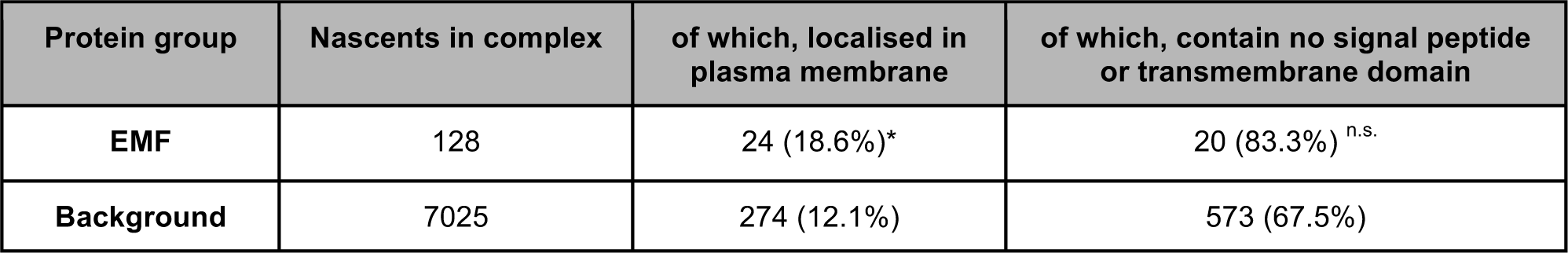
Numbers of nascent proteins localized in the plasma membrane and without conventional translocation signals, using Human Protein Atlas annotations. Percentages denote the proteins retained compared to the previous column. Where indicated, Fisher’s exact tests were performed to test for statistical significance using as background the set of 7025 nascent proteins liable to be assessed for 3’UTR-protein complexes. The number of proteins in this background that associated to the plasma membrane is 849, of which 573 lack a membrane addressing signal. The Benjamini-Hochberg procedure was applied for multiple test correction. Significance: ‘*’ indicates a P-value < 0.05; ‘**’ indicates a P-value < 0.01; ‘***’ indicates a P-value < 0.001; ‘n.s.’ indicates a P-value>0.05 and P-value<0.2.

**Supplementary Table 4.** Number of 3’UTR-protein complexes expected by chance (known moonlighting proteins). As for EMF proteins, the total number of 3’UTR-protein complexes expected by chance was measured on protein-protein interaction networks that had their protein labels randomly shuffled without replacement. Empirical p-values were calculated for the hypothesis that the number of 3’UTR-protein complexes formed using the real protein-protein interaction network is higher than the null distribution of 10,000 randomizations. For each moonlighting protein group, the number of predicted complexes, and the mean number of total complexes and its standard deviation (in parenthesis) over the randomizations is presented.

| Protein group | Nascent in complex | 3'UTR-protein complexes | 3'UTR-protein complexes in reshuffled networks | Empirical p-values |
| --- | --- | --- | --- | --- |
| **MoonProt (61)** | 32 (52%) | 163 (43) | 43.4 (25.1) | 0.0045 |
| **MoonDB (45)** | 15 (33,3%) | 81 (17) | 17.0 (10.9) | 0.0021 |
| **Both (88)** | 42 (47,7%) | 234 (55) | 55.4 (27.9) | 0.0017 |

